# Parental effects and the evolution of phenotypic memory

**DOI:** 10.1101/027441

**Authors:** Bram Kuijper, Rufus A. Johnstone

## Abstract

Despite growing evidence for nongenetic inheritance, the ecological conditions that favor the evolution of heritable parental or grandparental effects remain poorly understood. Here, we systematically explore the evolution of parental effects in a patch-structured population with locally changing environments. When selection favors the production of a mix of offspring types, this mix differs according to the parental phenotype, implying that parental effects are favored over selection for bet-hedging in which the mixture of offspring phenotypes produced does not depend on the parental phenotype. Positive parental effects (generating a positive correlation between parental and offspring phenotype) are favored in relatively stable habitats and when different types of local environment are roughly equally abundant, and can give rise to long-term parental inheritance of phenotypes. By contrast, unstable habitats can favor negative parental effects (generating a negative correlation between parental and offspring phenotype), and under these circumstances even slight asymmetries in the abundance of local environmental states select for marked asymmetries in transmission fidelity.

## 1 Introduction

Traditionally, evolution by natural selection relies on genetic inheritance. The possibility of nongenetic inheritance has, however, begun to attract increasing interest. Nongenetic inheritance refers to any effect of ancestors on descendants that is brought about by the transmission of factors other than sequences of DNA from parents or more remote ancestors (Bonduriansky & Day, 2009; Day & Bonduriansky, 2011; Jablonka & Raz, 2009; Danchin *et al*., 2011). Such effects may be mediated by a variety of different mechanisms, including the transmission of behaviour and culture by social learning (Richerson & Boyd, 2005; Aoki & Feldman, 2014), the transmission of epigenetic variants through heritable changes in DNA methylation or chromatin structure (Youngson & Whitelaw, 2008; Becker *et al*., 2011; Roux *et al*., 2011; Schmitz *et al*., 2011), the transmission of somatic or cytoplasmic factors such as hormones (Groothuis & Schwabl, 2008), nutrients or other maternal factors (Mousseau & Fox, 1998; Maestripieri & Mateo, 2009). All of these diverse mechanisms give rise to a trans-generational form of phenotypic plasticity, in which the phenotype of an offspring depends not only upon the genes it inherits and the environment to which it is exposed, but also upon the phenotype or environment of its parents or more remote ancestors (Uller, 2008; Smiseth *et al*., 2008; Shea *et al*., 2011).

There is a substantial amount of theoretical work that focuses on the consequences of nongenetic inheritance to other evolutionary processes (e.g., Kirkpatrick & Lande, 1989; Slatkin, 2009; Furrow *et al*., 2011; Day & Bonduriansky, 2011; Hoyle & Ezard, 2012; Townley & Ezard, 2013; Ezard *et al*., 2014). Yet, much less attention has been devoted to the evolution of such nongenetic inheritance mechanisms themselves, with the majority of studies focusing on specific mechanisms such as social learning (reviewed in Aoki & Feldman 2014). Only recently have models started to assess when and where nongenetic effects evolve (Leimar & McNamara, 2015; English *et al*., 2015; Kuijper & Hoyle, 2015). Interestingly, these studies have shown that parental cues on offspring phenotype determination are most likely to evolve when environments are correlated between parental and offspring generations (Kuijper & Hoyle, 2015), either because these environments are i) temporally stable (Leimar & McNamara, 2015; English *et al*., 2015) or fluctuate predictably (Kuijper & Hoyle, 2015); ii) have low dispersal between environments and iii) allow for transmission of cues between generations with little error (Leimar & McNamara, 2015; English *et al*., 2015).

Despite these important first insights, however, several questions about the evolution of nongenetic inheritance remain: first, it is currently poorly understood whether the evolution of these parental cues on phenotype determination will indeed result in *transmission* of phenotypes between generations, or whether other outcomes are possible. Often, nongenetic inheritance is associated with forms of ‘phenotypic memory’, signifying that phenotypes are stably transmitted from parent to offspring for a number of generations (Vastenhouw *et al*., 2006; Rando & Verstrepen, 2007; Becker *et al*., 2011; Rechavi, 2014). This requires a certain amount of parent-offspring resemblance and thus a positive correlation between parental and offspring phenotypes (often indicated as a positive parental effect: Räsänen & Kruuk, 2007; Kuijper *et al*., 2014; Kuijper & Hoyle, 2015). However, parent-offspring resemblance is not the only possible outcome, as parents may also induce offspring to attain a phenotype dissimilar to their own (often indicated as a negative parental effect; e.g., Janssen *et al*., 1988; Räsänen & Kruuk, 2007; Sikkink *et al*., 2014; Kuijper & Hoyle, 2015), which precludes the stable transmission of a phenotype across generations. In the current study, we aim to address not only when and where parental cues evolve, but also whether they result in phenotypic inheritance and of what form (i.e., duration of phenotypic memory).

Second, studies on phenotypic plasticity (Sultan & Spencer, 2002), local adaptation (reviewed in Kawecki & Ebert, 2004) and bet-hedging (Salathé *et al*., 2009; Gaál *et al*., 2010) find that asymmetries in the frequencies of different environments (e.g., combinations of very common and very rare environments) favor phenotypically monomorphic populations without plasticity or bet-hedging. How nongenetic inheritance is affected by asymmetric environments has, however, not been systematically addressed so far (English *et al*., 2015; Leimar & McNamara, 2015), particularly when considering environments that vary both at a spatial and temporal scale. Here we therefore assess the combined roles of spatial and temporal fluctuations on the evolution of nongenetic inheritance.

Third, existing models only consider large and well-mixed populations. However, another key insight from the bet-hedging literature is that populations with interacting relatives selectively favor more bet-hedging (and thus a larger amount of phenotypic variation) relative to well-mixed populations (Moran, 1992; Leimar, 2005; Gardner *et al*., 2007; Lehmann & Balloux, 2007). This raises the question whether interactions between relatives also affect the evolution of nongenetic inheritance, as its resulting patterns of inheritance affect the amount of phenotypic variation among offspring (e.g., Kuijper *et al*., 2014).

To address these three points, we use a model that is similar to that of Leimar & McNamara (2015) by considering development as a ‘switching device’, that integrates genetic and parental cues as inputs, and specifies offspring phenotypes as output. For the sake of clarity, we only focus on the evolution of parental cues relative to genetic cues, whereas environmental cues are considered in a follow-up paper. We treat phenotype as a binary variable, so that the developmental switching device determines the probability that an offspring is of one or the other of two alternative morphs. Similarly to Leimar & McNamara (2015), we suppose that the properties of this switching device may change over evolutionary time, as a result of the successive substitution of mutant modifier alleles that slightly alter its characteristics. These modifiers can be thought of as coding for the molecular machinery that mediates the transmission of a phenotypic effect from mothers to offspring, which can be most likely considered as a modifier that codes for a maternal hormone that affects the offspring’s phenotype (e.g., Groothuis & Schwabl, 2008; Gil, 2008), small RNAs (Ashe *et al*., 2012; Liebers *et al*., 2014) or DNA-methyltransferases (Lillycrop *et al*., 2007; Johannes *et al*., 2009).

## 2 The model

We focus on a sexual, ‘infinite island’ population (Wright, 1931, 1951), comprising infinitely many patches linked by juvenile dispersal. The assumption of infinitely many patches substantially simplifies the calculations of the frequencies of the different types, yet appears to be robust, as individual-based simulations that assume a finite number of patches (see below) give identical results. Each patch has a fixed population size of *n* breeders and switches back and forth between two possible environmental states **e** = {*e*_1_, *e*_2_}. Individuals can adopt one of two possible phenotypes **z** = {*z*_1_*, z*_2_}. Individuals are ‘locally adapted’ (and consequently experience a lower mortality rate) when their phenotype *z_i_* is identical to the environmental state *e_i_* of their patch. Within this fluctuating environment, individuals are characterized by a genetically-determined strategy (*p*_1_,*p*_2_), the elements of which specify the probability that an offspring will be of phenotype *z*_1_ when its parent is of phenotype *z*_1_ or *z*_2_, respectively. This strategy, which defines the rules of phenotypic transmission and inheritance, might (for instance) be encoded by genes coding for methylation proteins or other molecular machinery that regulates the non-genetic transmission of information between generations. A strategy for which *p*_1_ = *p*_2_ leads to the production of the same mix of offspring types regardless of the parental phenotype, implying that there is no transmission of phenotypic information from parent to offspring (or that such information, if transmitted, is not used). If selection favors a strategy for which *p*_1_ ≠ *p*_2_, by contrast, this implies that selection favours at least some influence of the parental phenotype on the determination of offspring phenotype, i.e. some degree of trans-generational plasticity. We use an adaptive dynamics approach (Geritz *et al*., 1998; McGill & Brown, 2007; Dercole & Rinaldi, 2008) to model the evolution of these strategies, assuming that evolutionary change occurs through the successive substitution of mutations of small effect. We thereby assume a separation of timescales: demographic changes (environmental change, deaths and births) occur at a much faster timescale than evolutionary change in the gene loci that specify the probabilities of phenotypic inheritance. We also allow for evolutionary change in the dispersal rate, *d*, on the same time scale as changes in the probabilities of phenotypic inheritance.

### Population dynamics

We model population dynamics in continuous time, such that demographic events (switches in the environmental state of a patch, deaths and births) occur sequentially, one-at-a-time (Moran, 1958; Metz & Gyllenberg, 2001; Alizon & Taylor, 2008). The following two types of event can occur:

i. *environmental change:* a single patch in environmental state *e_j_* switches to environmental state *e_i_* with rate 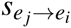 When the environment of a particular patch changes, all locally adapted individuals in the patch become locally maladapted, while all previously locally mal-adapted individuals become locally adapted. Note that 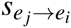 reflects the rate of change of an individual patch, rather than the global environment. Hence, at equilibrium, the total frequency of patches in one state versus the other is constant and given by 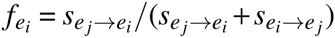
ii. *breeder mortality, birth and replacement:* the mortality rate of a *z_i_* breeder in an *e_j_* environment is given by *M_ij_*, where *M*_11_ < *M*_21_ and *M*_22_ < *M*_12_, implying that a breeder that is adapted to its current local environment has a lower mortality rate than a breeder that is maladapted to it. Whenever a death occurs, the now vacant breeding position is immediately filled (maintaining a local population size of *n* = 2), with probability *h* ≡ (1 − *d*) / (1 − *kd*) by the offspring of a random local breeder, and with probability 1 − *h* by the offspring of a random nonlocal breeder, where *d* denotes the juvenile dispersal rate and *k* the mortality cost of dispersal. The probability that a newborn juvenile is of phenotype *z*_1_ or *z*_2_ depends upon its parent’s phenotype through the strategy (*p*_1_, *p*_2_), as detailed above. Consequently, as deaths and births affect the distribution of various phenotypes over patches that differ in environmental quality, they also affect the reproductive values of each breeder and relatedness coefficients among breeders (see Supplementary Mathematica Sheet).

### Evolutionary dynamics

Let 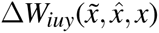 be the fitness effect of a focal mutant living in an environment *e_i_*-patch, who expresses a slightly deviant phenotype 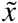, while its neighbour and the global population express phenotypes 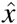 and *x* respectively. The subscript *iuy* reflects the state of the local environment (*e_i_*), whether the focal is currently adapted or maladapted to the local environment (*u ∈* {*a, m*}) and whether its neighbour is currently adapted or maladapted (*y ∈* {*a, m*}). In the Supplementary Information S1 we derive an expression for the fitness effect 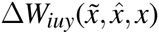 based on the mutant transition rates between the various classes of individuals. Note that these transition rates are weighed by the corresponding reproductive values as we consider a heterogeneous population (for similar approaches, see Alizon & Taylor, 2008; Taylor, 2009; Wild *et al*., 2009). By averaging over all possible environments i and individual states (*u,y*) and employing a standard result from inclusive fitness theory (Hamilton, 1964; Taylor & Frank, 1996), we obtain the following selection gradient Δ*W_x_* on phenotype *x* (see Supplementary Information S1)

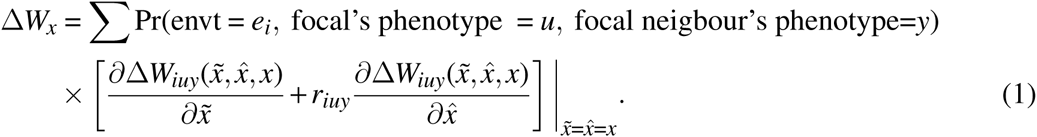

The first term in brackets reflects the fitness effect of a mutant allele borne by the focal breeder with phenotype 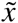, while the second term reflects the fitness effect of mutant allele borne by the focal’s neighbour (having phenotype 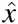), weighted by the coefficient of relatedness *r_iuy_* to the focal. Taking relatedness into account is essential in viscous populations, because the trait (*p*_1_, *p*_2_) affects local competition between relatives, as it changes the rate at which breeding positions become available in the local patch. To see this, note that the value of the trait (*p*_1_, *p*_2_) expressed by all relatives affects the probability that a maladapted versus adapted offspring will establish itself as a local breeder in the next timestep. In case this new breeder is adapted, it will occupy its breeding position for a longer and thus increases local competition by limiting the opportunities for offspring from other relatives to successfully establish themselves in the local patch.

If evolution proceeds slowly, so that an individual’s lifespan represents only an infinitesimal fraction of evolutionary time, then a standard result in adaptive dynamics (e.g., Dieckmann & Law, 1996; Day & Taylor, 2003) shows that one can describe the change in trait values over time with the continuous evolutionary time dynamic

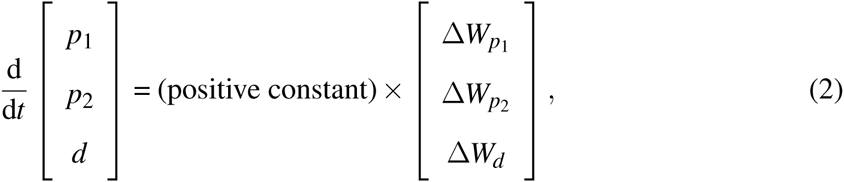

where we assume that pleiotropic mutations which simultaneously affect multiple traits are absent. We can use this dynamic to find singular strategies where the dynamic vanishes 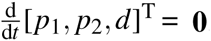 To find these strategies, we iterate the adaptive dynamic above in (2) starting from a particular set of values [*p*_1_, *p*_2_, *d* ]*_t_*_=0_ until values converged. During each time step of the iteration, we numerically solved for equilibrium values of patch type frequencies, reproductive values and relatedness for the current values of [*p*_1_, *p*_2_, *d* ]*_t_* using Euler’s method. Convergence was determined when the largest difference in values of [*p*_1_, *p*_2_, *d* ] between consecutive time steps was ≤ 10^−7^. Starting values used in our iterations are [*p*_1_, *p*_2_, *d* ]*_t_*_=0_ = [0.5,0.5,0.5]. The outcomes obtained from these numerical iterations are convergence stable by definition, and individual-based simulations revealed that values are also evolutionary stable, provided we assume that mutations occur independently in each strategic variable.

## 3 Results

In this section, we summarize how the evolutionarily stable strategy (*p*_1_, *p*_2_) varies with the parameters of the model (note that over the whole of the parameter range we consider, the model yields one unique evolutionarily stable outcome for any given set of parameter values). The key parameters that determine the nature of the environment are the switching rates 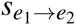 and 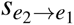. Rather than work directly with these switching rates, however, we characterize the environment in terms of two alternative, derived parameters: the relative frequency of local environment 2 compared to local environment 1, 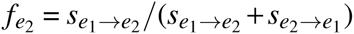 and the overall temporal instability of the environment, measured as the arithmetic average of the log switching rate across both patch states, 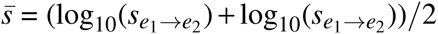 The former value matters because markedly asymmetric environments, in which one environmental state is much more common or exerts stronger selective pressures than the other, have been shown to substantially decrease the evolution of phenotype switching (Salathé *et al*., 2009; Gaál *et al*., 2010), relative to cases in which different environmental states have similar frequencies (e.g., Jablonka *et al*., 1995; Thattai & van Oudenaarden, 2004). Finally, we investigate the impact of the cost of dispersal, *k*.

In Figure 1 we plot the evolutionarily stable values of *p*_1_ and of *p*_2_ as a function of both 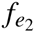 (frequency of local environment 2) and 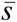 (mean log rate of switching between local environmental states), for several different values of *k* (the cost of dispersal). Corresponding to the values in Figure 1, Figure 2 depicts in a more schematic form the boundaries between those regions of parameter space in which *p*_1_ = *p*_2_ (implying that there is no transmission of phenotypic information from parent to offspring), those in which *p*_1_ > *p*_2_ (implying that there is a positive correlation between parental and offspring phenotypes), and those in which *p*_1_ < *p*_2_ (implying that there is a negative correlation between parental and offspring phenotypes). The evolutionarily stable values of *p*_1_ and of *p*_2_ determine the rules of phenotypic transmission and inheritance; in particular, they determine over how many generations on average each phenotype may be expected to persist before a switch occurs. In Figure 3 we plot the fidelities of inheritance at evolutionary equilibrium, i.e. the expected number of generations over which each phenotype is faithfully copied, again as a function of both 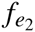 (frequency of local environment 2) and 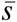 (mean log rate of switching between local environmental states), for several different values of *k* (the cost of dispersal).

**Figure 1.**
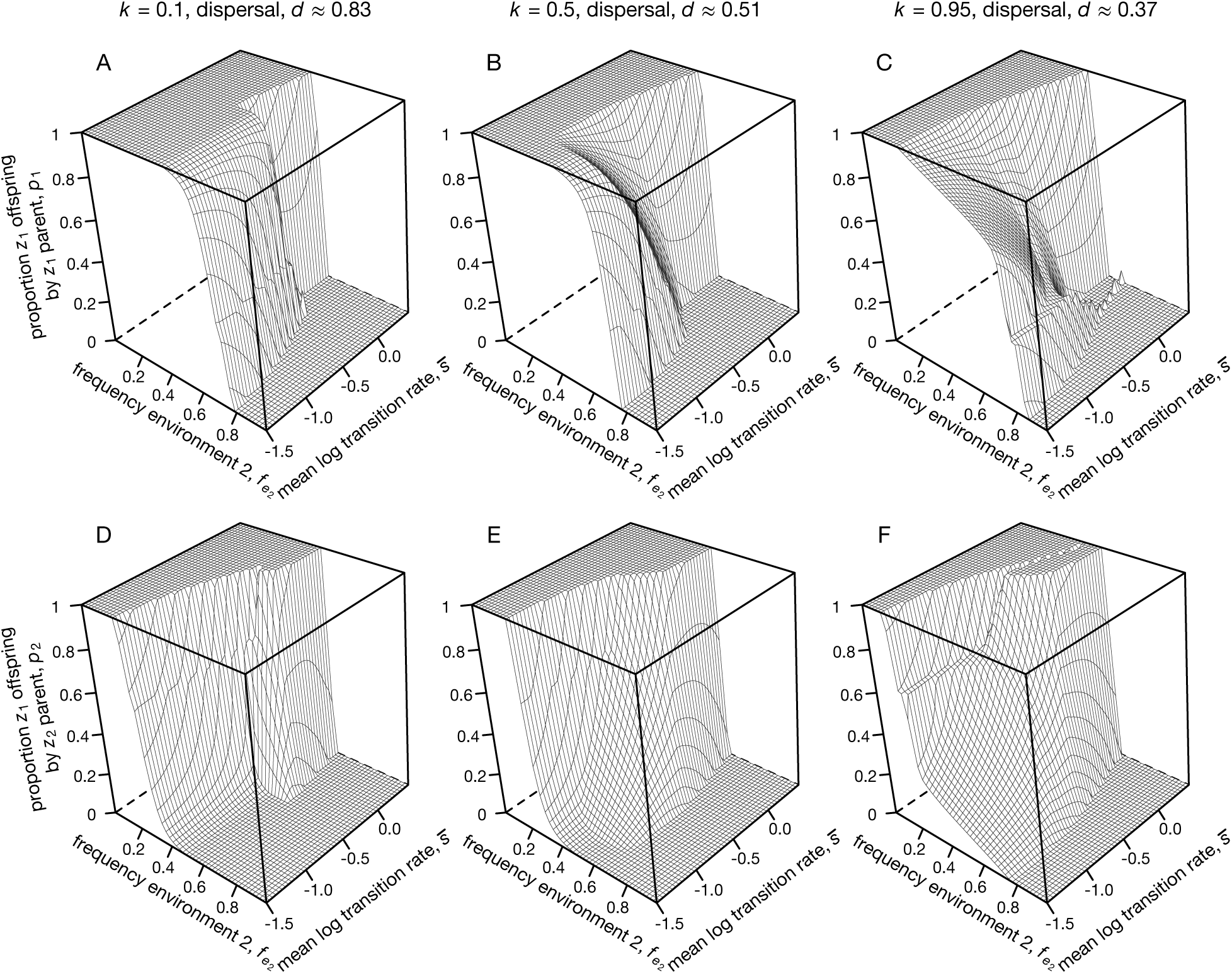
Equilibrium values of *p*_1_ (panels A-C) and *p*_2_ (panels D-F), which reflect the equilibrium proportions of *z*_1_-offspring produced by parents with phenotypes *z*_1_ and *z*_2_ respectively. In each panel, we explore equilibrium outcomes over a range of different parameter values: along the *x*-axis, we vary the relative frequency of environment 2, 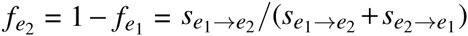 along the *y*-axis, we vary environmental stability, measured as the average log_10_ environmental switch rate 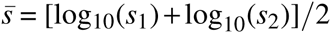 Parameters: mortality rates maladapted versus adapted = 2:1.

**Figure 2.**
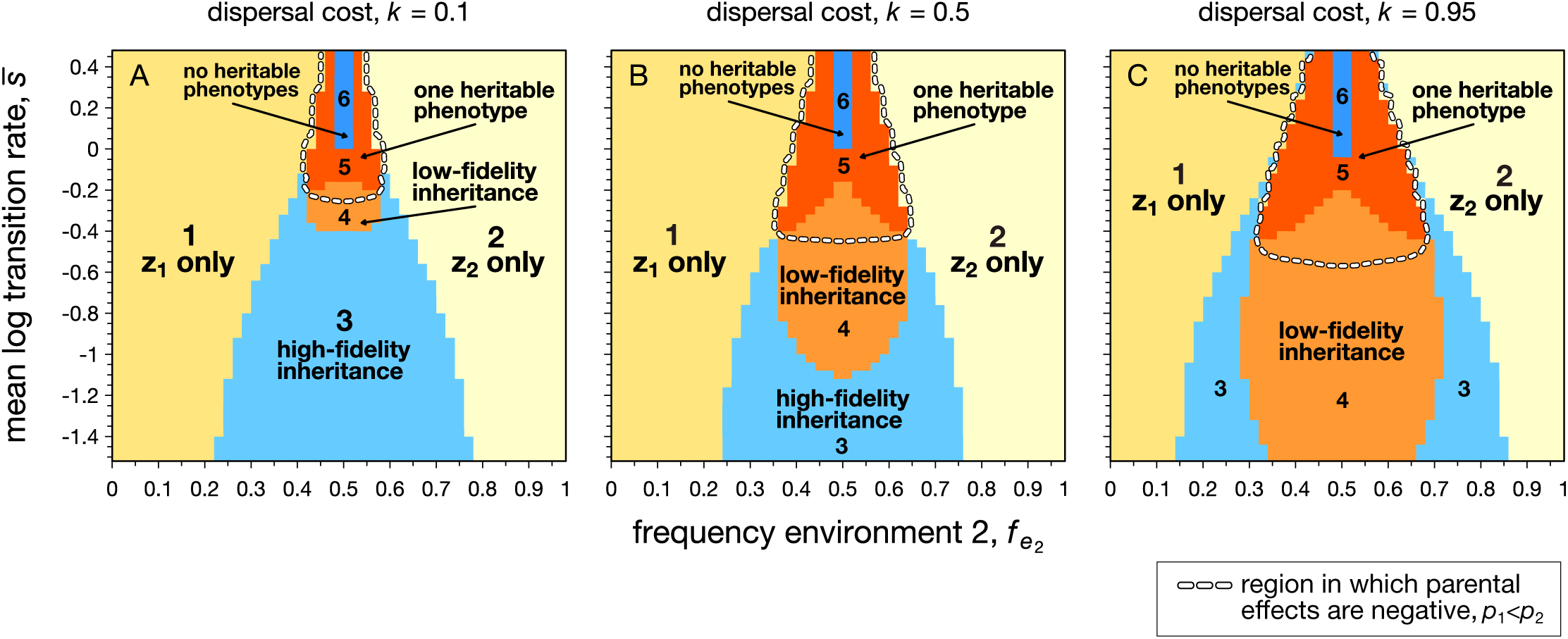
A summary of the overall fidelity of inheritance that results from the evolved values of *p*_1_ and *p*_2_ as displayed in Figure 1. *Region 1: p*_1_ = *p*_2_ = 1, only phenotype *z*_1_ persists in the population. *Region 2: p*_1_ = *p*_2_ = 0, only phenotype *z*_2_ persists in the population. *Region 3:* high-fidelity inheritance of both phenotypes: *p*_1_ = 1, 0 ≤ *p*_2_ < 1 or 0 < *p*_1_ ≤ 1, *p*_2_ = 0. Both phenotypes persist in the population, but are copied with high-fidelity (i.e., at least one of the two parental phenotypes transmits its phenotype always to all offspring). *Region 4:* low-fidelity inheritance, 0 < *p*_1_, *p*_2_ < 1, as neither parental phenotype transmits its phenotype with the highest fidelity to offspring. *Region 5:* only one phenotype is heritable, whereas the other phenotype will be produced by parents of the opposite phenotype. *p*_1_ = 0, 0 < *p*_2_ < 1 or *p*_2_ = 1, 0 < *p*_1_ < 1. *Region 6:* neither phenotype is heritable from parent to offspring; parents always produce offspring with phenotypes opposite to their own. *p*_1_ = 0, *p*_2_ = 1.

**Figure 3.**
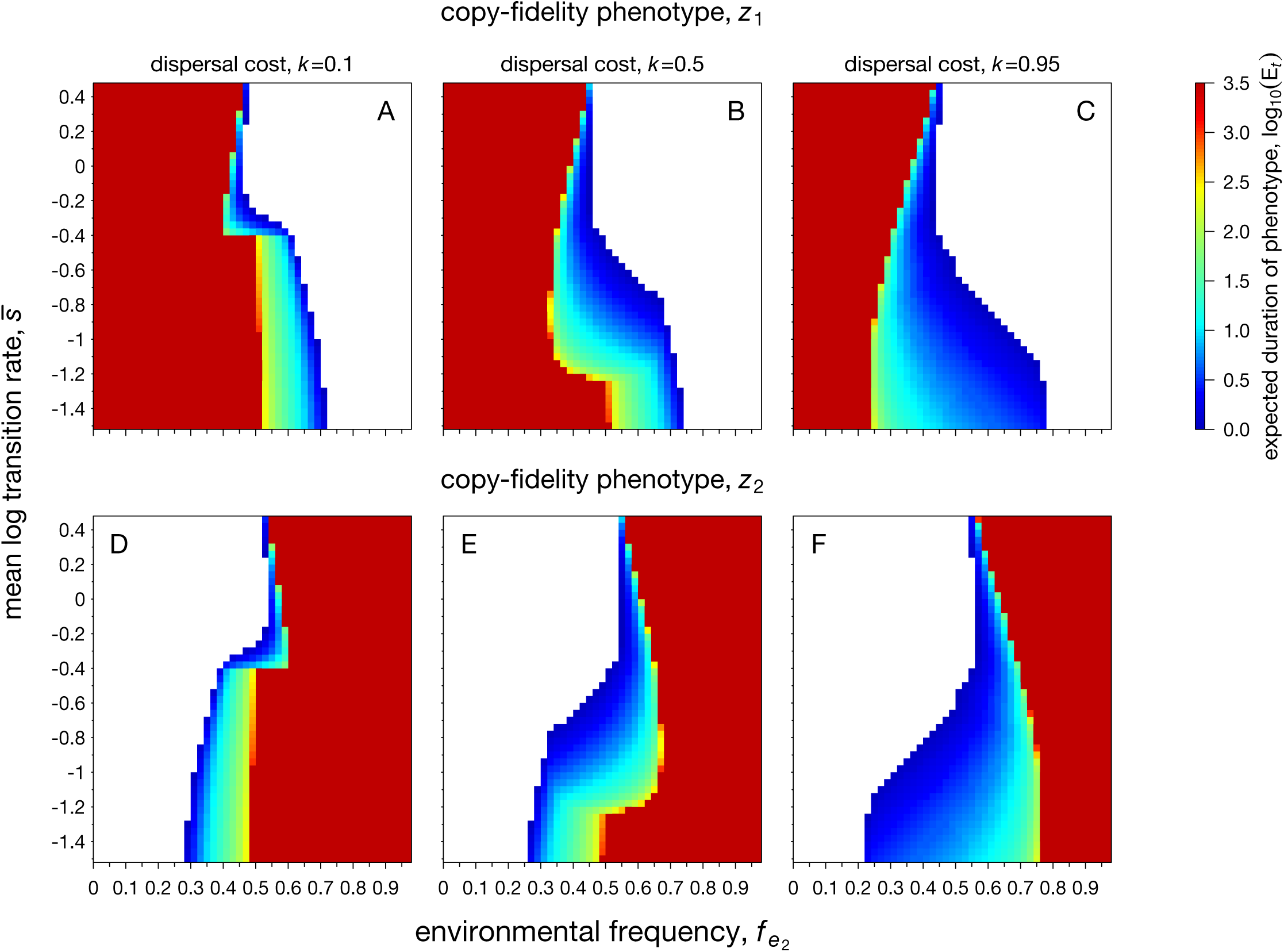
Heat plots depicting the expected number of generations *E_t_* that a phenotype is copied from parent to offspring. Panels A-C: expected number of generations that phenotype *z*_1_ is copied to descendants. Panels D-F: expected number of generations that phenotype *z*_2_ is copied is copied to descendants. Deep blue indicates that a particular phenotype lasts only a single generation. White indicates absence of nongenetic inheritance. Parameters as in Figure 1.

Below, we consider the effects of these parameters and summarize (with reference to the figures) their impact on the form of the evolutionarily stable strategy.

### Asymmetries in the frequencies of local environments

When one environment is much more common than the other, Figures 1 and 2 show that the evolutionary outcome is a monomorphism. Both parental phenotypes produce only *z*_1_-offspring (*p*_1_ = *p*_2_ = 1) when environment 1 is common, or only *z*_2_-offspring (*p*_1_ = *p*_2_ = 0) when environment 2 is common. Under these circumstances, selection favours insensitivity to parental phenotype on the part of developing young because the majority of offspring are likely to encounter only the most common environment, regardless of their parent’s phenotype. These outcomes resemble a scenario of genetic inheritance with extremely low mutation rates. Figure 3 shows that when environment 1 is common, phenotype *z*_1_ is copied with very high fidelity while phenotype *z*_2_is almost never transmitted from parent to offspring, and similarly when environment 2 is common, phenotype *z*_2_ is copied with very high fidelity while phenotype *z*_1_is almost never transmitted.

It is only when different environments are encountered at more similar rates that the model yields a phenotypically polymorphic outcome. Under these circumstances, selection almost always favours some degree of sensitivity to parental phenotype on the part of developing young, with *p*_1_ diverging from *p*_2_ (see Figure 1). The precise values of *p*_1_ and *p*_2_ depend strongly upon the degree of environmental stability, and on the cost of dispersal, as discussed below. Nevertheless, even before we consider the impact of these parameters, it is clear that strict bet-hedging, in which parents produce a mixture of the two offspring types that is independent of their own phenotype (De Jong *et al*., 2011), is almost never favoured.

### Environmental stability

Focusing in more detail on the evolutionarily stable values of *p*_1_ and *p*_2_ at a polymorphic equilibrium, Figure 2 reveals that more temporally stable environments (characterized by lower values of 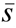) favour more faithful transmission of parental phenotypes, i.e. they lead to outcomes at which *p*_1_ ≫ *p*_2_ (so that a *z*_1_-parent is very likely to produce *z*_1_-offspring, and a *z*_2_-parent to produce *z*_2_-offspring). The reason is that, when patches rarely switch state, the lower mortality of locally adapted individuals gives rise to a strong positive correlation between an individual’s phenotype and the state of its local environment. Under these circumstances, philopatric offspring are likely to experience an environment matching their parent’s phenotype, and so they benefit by adopting the same phenotype themselves. Of course, it is only philopatric offspring that are likely to benefit from transmission of phenotypic information in this way; offspring that disperse encounter an environment that is uncorrelated with their natal environment, and hence with their parent’s phenotype. However, a positive correlation between parental phenotype and offspring environment for philopatric young, and an absence of any correlation for dispersing young, yields a correlation that is positive overall.

As the rate of environmental switching 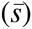 increases, the correlation between parental phenotype and environmental state weakens and selection favors less faithful transmission of phenotypes (i.e. the values of *p*_1_ and *p*_2_ converge). This reflects the action of kin selection – limited dispersal leads to local relatedness among breeders in a patch, with philopatric offspring breeding alongside their parents. As a result, breeders are selected to produce a fraction of offspring who differ from them in phenotype, so that in the (rare) event of environmental change, at least one of parent or offspring are adapted to the new environment, securing the long-term survival of their genotype. Nevertheless, the precise mix of offspring types reflects the parental phenotype, which predominates over the alternative phenotype. Parents, in other words, produce a majority of offspring matching their own phenotype, and only a minority of the alternative phenotype. Since this leads to a positive correlation between parent and offspring phenotypes (*p*_1_ > *p*_2_), we consider it an example of a positive parental effect. Additionally, such positive parental effects can also be considered an example of condition-dependent bet-hedging sensu De Jong *et al*. (2011), since mixtures of young are dependent on the parental phenotype.

As the rate of environmental switching increases still further, selection favors still greater rates of phenotype switching, to the extent that the correlation between parental and offspring phenotypes eventually changes sign and becomes negative (*p*_1_ < *p*_2_). Parents, in other words, transmit their own phenotype only to a minority of their offspring, while the majority are of the opposite phenotype, an example of a negative parental effect. It is worth emphasizing that even under these circumstances, the lower mortality rate of locally adapted individuals still maintains a positive (though weak) correlation between an individual’s phenotype and the state of its local environment. Why then should selection favor negative parental effects under these circumstances? Since individuals that are maladapted to their local environment have a higher death rate, more empty breeding positions are available in patches that contain maladapted individuals. In turn, this makes it more likely that philopatric offspring from a maladapted parent obtain a breeding position as opposed to philopatric offspring from an adapted parent. As a result, despite the overall positive correlation between phenotype and environment across the adult population, there is a negative correlation between the phenotype of the parent of a newly established breeder and the environment that breeder experiences. This negative correlation favors the evolution of negative parental effects.

Figure 3 shows the consequences of these changes in *p*_1_ and *p*_2_ for the fidelity of inheritance. When both environments are similarly common (around 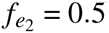), more stable environments favour longer-term phenotypic memory, with phenotypes copied across a larger number of generations on average, while less stable environments favour shorter-term memory. In the extreme, when there is a negative correlation between parental and offspring phenotype (*p*_1_ < *p*_2_), switching becomes more likely than faithful transmission, and the average duration of a phenotype drops below 2 generations.

### Interaction between asymmetrical frequencies of local environments and environmental stability

As we have seen, as environments change from stable to unstable there is a shift from more faithful to less faithful transmission of phenotypes, and thus from positive to negative parental effects. When 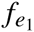 is equal to one half, implying that both environments are equally common, this shift happens in parallel for both phenotypes, which are always transmitted with equal degrees of fidelity. But as the relative abundance of the two environmental states deviates from a 1:1 ratio, asymmetries in transmission are favored, with the phenotype matching the more common environment being transmitted more faithfully than the phenotype matching the less common environment. Further, there is an interaction between environmental stability and the relative abundance of environmental states, such that asymmetries in transmission evolve more readily in unstable environments.

When patches change state only rarely, even quite substantial differences in the abundance of the two environments favor no or minor differences in the fidelity of transmission of the two corresponding phenotypes. By contrast, when environments change frequently, even slight differences in the relative abundance of the two environments favors strongly asymmetrical transmission. The region of negative parental effects bounded by the dotted line (which is associated with rapid environmental switching) is thus dominated by zone 5, in which only one phenotype can be transmitted from parent to offspring: on the left part of zone 5, for instance, environment *e*_1_ is more common than *e*_2_, and the parental phenotype that matches the common environment, *z*_1_, produces a mixture of both phenotypes, whereas the phenotype matching the rarer environment, *z*_2_, will exclusively produce offspring of the opposite (common) type. The reverse applies to the right part of region 5, with *z*_1_ parents exclusively producing offspring of the opposite phenotype, *z*_2_ (*p*_1_ → 0) and *z*_2_ parents producing offspring with their own *z*_2_ phenotype as well as offspring with the opposite phenotype *z*_1_ (0 < *p*_2_ < 1). In essence, region 5 implies that only one phenotype is directly transmitted from parent to offspring (the phenotype matching the common environment), whereas the other phenotype (fitting to the rare environment) originates solely from parents that do not express that phenotype themselves. Such a scenario resembles certain cases of bacterial persistence (Gardner *et al*., 2007; Lewis, 2010), in which a normal phenotype gives rise to both normal offspring and occasionally to offspring with a persister phenotype, whereas persister phenotypes almost exclusively give rise to offspring with a normal phenotype that is different to their own.

As the relative abundance of the two environmental states deviates from a 1:1 ratio, asymmetries in phenotype transmission are favored, with the phenotype matching the more common environment being transmitted more faithfully than the phenotype matching the less common environment (see Figure 3). Furthermore, there is an interaction between environmental stability and the relative abundance of environmental states, such that asymmetries in transmission evolve more readily in unstable environments.

### Limited dispersal and interactions with relatives

Comparing the three columns of Figure 1, we see that greater costs of dispersal, leading to a higher frequency of philopatry at equilibrium and thus to greater local relatedness promotes the divergence of *p*_1_ from *p*_2_ across a wider range of the parameter space. Moreover, Figure 3 shows that such costs tend to promote less faithful transmission of parental phenotypes, i.e., they lead to less positive or to more negative parental effects. This reflects the action of kin selection: diversification of young enhances the longterm survival of the local genotype by ensuring that at least one of the carriers of this genotype will be adapted after any future environmental change. Since reduced dispersal entails a higher degree of relatedness among local breeders, it promotes behavior that is beneficial at the level of the local group, and therefore favors less faithful inheritance, since this acts to create local phenotypic diversity among carriers of the genotype in question. Figures 3C,F indeed show that reduced dispersal in combination with modestly stable environments leads to a broad range of intermediate inheritance fidelities that are congruent with nongenetic inheritance (summarized in Figure 2C).

## 4 Discussion

Similar to previous studies (Rivoire & Leibler, 2014; English *et al*., 2015; Leimar & McNamara, 2015; Kuijper & Hoyle, 2015), our model shows that selection can favour mechanisms of phenotype determination in offspring that are sensitive to parental phenotype as a cue, whenever the parental phenotype is correlated with the environment that offspring are likely to encounter. Depending upon the sign of this correlation, selection can favour either positive or negative parental effects, such that offspring resemble their parent more or less than would be expected by chance (see Figure 2;). It may seem surprising that selection can so readily favour negative parental effects, given that selection invariably promotes a positive correlation between individual phenotypes and the state of their local environment. But local competition for breeding vacancies implies that offspring may have to wait for the death of their parent to claim such a vacancy, and this is more likely if the parent is locally maladapted. Hence, if the population-wide positive correlation between phenotypes and local environments is weak enough, offspring that are able to claim a breeding vacancy in their natal patch may experience an environment that is negatively correlated with their parent’s phenotype. As found in previous models (Kuijper *et al*., 2014; Kuijper & Hoyle, 2015), such a negative correlation is more likely to be found in an unstable habitat, where patches change state frequently.

Building on previous analyses of the evolution of parental cues (Rivoire & Leibler, 2014; Kuijper *et al*., 2014; Leimar & McNamara, 2015; Kuijper & Hoyle, 2015; English *et al*., 2015), we highlight three novel findings of the current analysis: first, we have found that the evolution of parental cues does not necessarily lead to inheritance (i.e., a resemblance between parental and offspring phenotypes). When negative parental effects evolve (see Figure 2), parents are more likely to produce offspring with a phenotype opposite to their own. In asymmetrical environments, such negative parental effects can even lead to substantial between-phenotype differences in inheritance fidelity (see Figure 3), where parents with a phenotype matching the rare environment never give rise to offspring with the same phenotype, whereas parents of the common phenotype produce a mixture of both phenotypes. In the presence of positive parental effects, however, we find that parental cues are likely to lead to nongenetic inheritance, lasting > 1 to several hundreds of generations (see Figure 3). To conclude, our model predicts that the inheritance fidelity resulting from nongenetic inheritance is not a fixed entity, but can be highly sensitive to the particular configuration of the environment, the level of dispersal and the phenotype in question.

Second, our model shows that asymmetries in the frequencies of both environments have a profound effect on the evolution of parental cues. Parental cues evolve in even or modestly asymmetric environments, but are disfavored in highly asymmetric environments (where 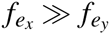) where a monomorphic population evolves instead. In the latter case, long-term selection on genotypes (most often occurring in the common environment) is sufficient to predict the future environment (Shea *et al*., 2011), so that information based on parental cues provides little additional information. Interestingly, selection against parental cues is exacerbated in rapidly changing environments (e.g., compare top and bottom of Figure 2), where smaller asymmetries can already result in a monomorphism. This is because in rapidly changing, asymmetric environments, individuals are certain to experience environmental change during their lifetime, so that all individuals live in the common environment during a part of their life. As a result, between-individual differences in selective conditions become smaller relative to slowly changing environments, reducing the selective need for parental information about the future. To summarize, parental cues should be most prevalent in contexts where different environments can be encountered at roughly similar rates.

A third novel aspect of our model is that it focuses on a spatially structured population in which relatives interact. We find that spatial structure and limited dispersal substantially influence the evolution of nongenetic inheritance, because they give rise to a higher degree of relatedness among local breeders, thereby favoring diversification of young to enhance the long-term survival of the group (Moran, 1992; Leimar, 2005; Lehmann & Balloux, 2007). This effect is strengthened by our assumption of overlapping generations, which ensures that philopatric offspring often coexist alongside their parents, creating a strong selective pressure for dissimilarity between parent and young. Our model thus predicts that parental effects should be more common in populations with interacting relatives relative to well-mixed populations. As parental effects receive a growing amount of attention in eusocial (reviewed in Linksvayer & Wade, 2005; Yan *et al*., 2014) and subsocial insects, it would therefore be timely and interesting to compare parental effects (e.g., heritable epimutations, heritable small RNAs) across related species that vary in the extent of their social interactions.

The impact of demography and interacting relatives on the evolution of parental cues offers many possibilities for further study. An example of a demographical process which deserves further attention in the context of parental cues is the ‘multiplier effect’ (McNamara & Dall, 2011; Dall *et al*., 2015). The multiplier effect occurs when cue-ignoring phenotypes do poorly in wrong environments, yet leave many copies of themselves in the right environment. As a consequence, particular genotypes accumulate in one environment only, diminishing selection for (parental) cues on phenotype determination. In our model, however, the presence of overlapping generations in our model reduces the multiplier effect: this is because successful parents live longer and thus occupy breeding positions for longer, so that the only offspring genotypes that are successful will be those that have migrated to remote environments (in which the environment is not necessarily the same). More importantly, the evolution of parental cues in our model does not directly limit the distribution of genotypes to certain environments, whereas the evolution of natal philopatry in McNamara & Dall (2011) directly restricts the range of environments seen by a genotype. That said, future studies should study the importance of the multiplier effect further, particularly in contexts i) in which generations are overlapping versus discrete, ii) when parental effects may restrict the range of environments experienced by offspring genotypes via other means than philopatry (e.g., by modulating the offspring’s social environment). Above all, the current model and that of McNamara & Dall (2011) clearly illustrate that demographical processes cannot be ignored when studying parental cues on offspring phenotype determination.

Our analysis is not restricted to any one particular mechanism by which cues about the parental phenotype might be transmitted to offspring. For example, the finding that stable environments select for positive parental effects and hence, transmission of phenotypes over a substantial number of generations, could potentially pertain to long-term transmission of epigenetic variants in *Arabidopsis thaliana* (Becker *et al*., 2011; Schmitz *et al*., 2011; Van der Graaf *et al*., 2015) or the self-reinforced transmission of parental care in rats, which is only disrupted when parents were experimentally induced to provide less care than normal (Kappeler & Meaney, 2010). Examples of negative parental effects have been found in nature, such as bacterial persistence, in which normal cells always give rise to a small minority of persister cells that are tolerant to antibiotics, whereas persister cells mainly produce offspring cells of a normal phenotype (Keren *et al*., 2004; Balaban *et al*., 2004; Lewis, 2010). Other cases in which parents negatively affect the phenotype of their offspring have been found in the context of negative maternal effects (reviewed in Räsänen & Kruuk, 2007), such as negative maternal influences on offspring age at maturity in springtails (Janssen *et al*., 1988), heat stress in *C. elegans* (Sikkink *et al*., 2014) or juvenile growth rates in red squirrels (McAdam & Boutin, 2003). Given our prediction that such positive and negative parental effects are connected to slow and rapidly changing conditions, assessing the role of environmental change in the examples above may provide key insights into the evolution of nongenetic effects.

An aspect that is beyond the scope of the current study are the long-term consequences of parental effects for phenotypic evolution. The major evolutionary consequence of the cooccurrence of nongenetic and genetic inheritance is a decoupling of the change in the mean phenotype from that of the mean genotype. This decoupling can lead to interesting dynamical consequences, such as non-equilibrium dynamics (Benton *et al*., 2001; Van Cleve & Feldman, 2008; Inchausti & Ginzburg, 2009), evolutionary momentum (Feldman & Cavalli-Sforza, 1976; Kirkpatrick & Lande, 1989; Lande & Kirkpatrick, 1990) and rapid adaptation combined with transient dynamics (Hoyle & Ezard, 2012; Townley & Ezard, 2013). Moreover, nongenetic inheritance can influence the intensity and the sign of selection acting on genetically inherited traits in unexpected ways (Bonduriansky *et al*., 2011), which has been exemplified by the substantial body of work on gene-culture coevolution (Feldman & Laland, 1996; Laland *et al*., 2010). Understandably, models exploring these effects typically assume that nongenetic effects exist a priori in order to investigate their consequences. Our focus, by contrast, has been to explore the circumstances that favor the evolution of parental effects. Only by integrating the evolution of parental effects and their consequences can we hope obtain a more complete picture about the role of parental effects as capacitors of evolutionary change (Badyaev, 2008).

## Acknowledgements

Jonathan Wells, Rebecca Hoyle, Stuart Townley and Thomas Ezard are thanked for valuable discussions within the Transgen group. This work was supported by EPSRC sandpit grant EP/H031928/1 (RAJ) and EPSRC grant EP/I017909/1 awarded to the 2020 Science consortium, which provided a fellowship to BK. The authors acknowledge the use of the UCL Legion High Performance Computing Facility (Legion@UCL), and associated support services, in the completion of this work. We thank the Lorentz Centre in Leiden, the Netherlands and the Dutch Royal Academy of Arts and Sciences (KNAW) for funding of a workshop on nongenetic effects that contributed to the results presented in this paper. The authors have no conflict of interest to declare.

